# USP15 participates in HCV propagation through the regulation of viral RNA translation and lipid droplet formation

**DOI:** 10.1101/432930

**Authors:** Shinji Kusakabe, Tatsuya Suzuki, Yukari Sugiyama, Saori Haga, Kanako Horike, Makoto Tokunaga, Junki Hirano, Zhang He, David Virya Chen, Hanako Ishiga, Yasumasa Komoda, Chikako Ono, Takasuke Fukuhara, Masahiro Yamamoto, Masahito Ikawa, Takashi Satoh, Shizuo Akira, Tomohisa Tanaka, Kohji Moriishi, Moto Fukai, Akinobu Taketomi, Sachiyo Yoshio, Tatsuya Kanto, Tetsuro Suzuki, Toru Okamoto, Yoshiharu Matsuura

## Abstract

Hepatitis C virus (HCV) utilizes cellular factors for an efficient propagation. Ubiquitin is covalently conjugated to the substrate to alter its stability or to modulate signal transduction. In this study, we examined the importance of ubiquitination for HCV propagation. We found that inhibition of de-ubiquitinating enzymes (DUBs) or overexpression of non-specific DUBs impaired HCV replication, suggesting that ubiquitination regulates HCV replication. To identify specific DUBs involved in HCV propagation, we set up an RNAi screening against DUBs and successfully identified ubiquitin-specific protease 15 (USP15) as a novel host factor for HCV propagation. Our studies showed that USP15 is involved in translation of HCV RNA and production of infectious HCV particles. In addition, deficiency of USP15 in human hepatic cell lines (Huh7 and Hep3B/miR122 cells) but not in a non-hepatic cell line (293T cells) impaired HCV propagation, suggesting that USP15 participates in HCV propagation through the regulation of hepatocyte-specific functions. Moreover, we showed that loss of USP15 had no effect on innate immune responses *in vitro* and *in vivo*. We also found that USP15-deficient Huh7 cells showed reductions in the sizes and numbers of lipid droplets (LDs), and addition of palmitic acids restored the production of infectious HCV particles. Taken together, these data suggest that USP15 participates in HCV propagation by regulating the translation of HCV RNA and formation of LDs.

## Importance

Although ubiquitination has been shown to play important roles in the HCV life cycle, the roles of de-ubiquitinating enzymes (DUBs), which cleave ubiquitin chains from their substrates, in HCV propagation have not been investigated. Here, we identified USP15 as a DUB regulating HCV propagation. USP15 showed no interaction with viral proteins and no participation in innate immune responses. Deficiency of USP15 in Huh7 cells resulted in suppression of the translation of HCV RNA and reduction in the sizes and amounts of lipid droplets, and addition of fatty acids partially restored the production of infectious HCV particles. These data suggest that USP15 participates in HCV propagation in hepatic cells through the regulation of viral RNA translation and lipid metabolism.

## Introduction

Hepatitis C virus (HCV) belongs to the *Flaviviridae* family and possesses a single-stranded positive-sense RNA as a genome (1). Viral RNA is translated to a precursor polyprotein which is cleaved into 10 viral proteins by host and viral proteases. Among the HCV proteins, core, E1 and E2 proteins form viral particles, and non-structural (NS) 3, 4A, 4B, 5A and 5B proteins are responsible for HCV RNA replication. NS2 protein cleaves the junction between NS2 and NS3, and p7 has been shown to exhibit ion channel activity (1). HCV infection leads to chronic infection and eventually induces steatosis, cirrhosis and hepatocellular carcinoma (2). HCV core protein localizes with many cellular components, such as nucleus, endoplasmic reticulum (ER), lipid droplets (LDs), lipid rafts and mitochondria (3–7). On the other hand, HCV infection epidemiologically correlates with extra-hepatic manifestations (EHMs) such as type 2 diabetes, mixed cryoglobulinemia and non-Hodgkin lymphoma (8). Liver-specific HCV core transgenic (CoreTG) mice develop insulin resistance, steatosis and hepatocellular carcinoma (9, 10), suggesting that HCV core protein plays a role in liver diseases and EHMs.

Efficient propagation of HCV requires several cellular factors, such as miR-122, a liver-specific microRNA that binds to two sites of HCV RNA to facilitate HCV replication (11, 12), and protein complexes of molecular chaperones and co-chaperones such as heat shock proteins, cyclophilin A, FK506-binding protein (FKBP) 8 and FKBP6 (13–15). In addition, phosphatidylinositol-4-kinase alpha/beta-mediated phosphatidylinositol-4-phosphate is required to construct appropriate membrane structure for HCV replication (16–18), and components of lipoproteins such as apolipoprotein (APO) E and APOB play important roles in the maturation of HCV particles (19–21). Lipid rafts, LDs and their associated proteins are also involved in HCV replication (22–24). Therefore, HCV utilizes various cellular organelles and host factors to facilitate an efficient propagation.

Ubiquitination is a post-translational modification that regulates cellular homeostasis. The HCV core protein was reported to be ubiquitinated by E6-associated protein (E6AP) to suppress viral particle formation (25). Blockage of the cleavage of core protein by signal peptide peptidase (SPP) has been shown to induce ubiquitination of core protein by translocation in renal carcinoma on chromosome 8 (TRC8) to suppress induction of ER stress in culture cells (26). Zinc mesoporphyrin (ZnMP) has been reported to induce degradation of NA5A via ubiquitination (27). It was also reported that interferon-stimulated gene (ISG)-12a (ISG12) induced by HCV infection ubiquitinates and degrades NS5A by S-phase kinase-associated protein 2 (SKP2) (28). NS5B was shown to interact with human homolog 1 of protein linking integrin-associated protein and cytoskeleton (hPLICs) to promote proteasomal degradation (29). In addition, HCV infection has been shown to induce ubiquitination of Parkin to promote mitophagy (30, 31) and regulate the ubiquitination of retinoic acid-inducible gene-I (RIG-I) through the ISG15/PKR pathway (32). These data suggest that ubiquitination participates in various steps of the HCV lifecycle.

In this study, we found that treatment with an inhibitor of de-ubiquitinating enzymes (DUBs) or overexpression of non-specific DUBs impaired HCV replication, suggesting that ubiquitination is important for HCV propagation. An RNAi-mediated screening targeting DUB genes identified ubiquitin-specific protease 15 (USP15) as a novel host factor that participates in HCV replication. Translation of HCV RNA was significantly impaired in USP15-deficient Huh7 cells (USP15KOHuh7). Deficiency of USP15 in hepatic but not in non-hepatic cell lines significantly reduced the propagation of HCV. Unlike in previous reports, we found that USP15 was not involved in RIG-I-mediated innate immune responses *in vitro* and *in vivo*. In addition, we found that the expression of sterol regulatory element-binding protein (SREBP)-1c, a master regulator of fatty acid synthesis and LDs, was suppressed in USP15KOHuh7 cells. USP15 was localized on LDs, and the addition of fatty acids restored the production of infectious HCV particles in USP15KOHuh7 cells. Taken together, these data suggest that USP15 is a crucial host factor for HCV propagation in hepatic cells through the regulation of viral RNA translation and lipid metabolism.

## Results

### Ubiquitination is required for HCV propagation

To examine the importance of ubiquitination during HCV replication, HCV replicon cells were treated with PR-619, a non-specific DUB inhibitor. Intracellular HCV RNA was significantly reduced by the treatment with PR-619 without obvious cytotoxicity (Fig. 1A), suggesting that inhibition of DUBs impaired HCV replication. OTUD7B, OTUB1 and OTUD1 are DUBs which specifically cleave the K11-, K48- and K63-linked ubiquitin chains from non-specific target proteins, respectively (33–35). Immunofluorescence microscopic observation of Huh7 cells overexpressing OTUD7B, OTUB1 and OTUD1 revealed the cytoplasmic localization of these DUBs, which suggested that all DUBs cleave ubiquitin chains of the target proteins in the cytoplasm (Fig. 1B and 1C). To examine which types of ubiquitination are involved in HCV replication, Huh7 cells overexpressing DUBs were infected with HCV. Overexpression of all DUBs in Huh7 cells impaired HCV propagation (Fig. 1D), suggesting that the K11-, K48- and K63-linked of ubiquitination are required for HCV propagation.

**Figure 1.**
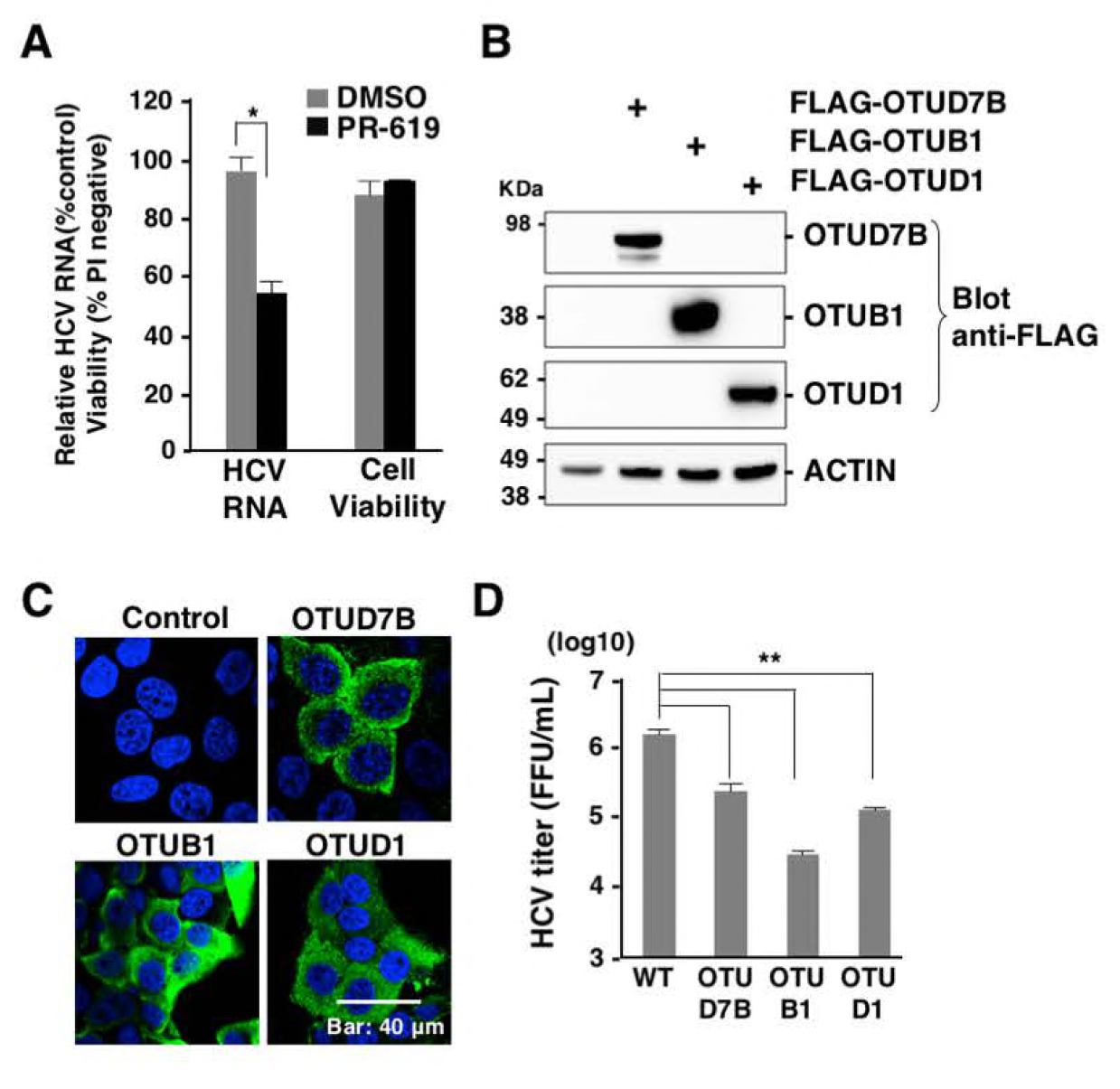
Ubiquitination is important for HCV replication. (A) HCV replicon (9–13) cells were treated with PR-619, a non-selective DUB inhibitor, or DMSO for 24 h, and cell viability and intracellular HCV RNA were determined by PI staining and qPCR, respectively. (B) Expression of FLAG-tagged OTUD7B, OTUB1 and OTUD1 in Huh7 cells was detected by Western blotting using anti-FLAG antibody. (C) Subcellular localization of OTUD7B, OTUB1 or OTUD1 overexpression in Huh7 cells was observed by confocal microscopy. Each DUB (green) or nucleus (blue) was stained with anti-FLAG antibody and DAPI, respectively. (D) HCV was infected into Huh7 cells expressing the indicated DUBs at an moi of 3. After 4 days post-infection, the culture supernatants were collected and infectious HCV titers in the culture supernatants were determined by a focus forming assay.

### RNAi screening to determine DUBs involved in HCV propagation

Because treatment with a DUB inhibitor and overexpression of non-specific DUBs suppressed HCV replication, we next tried to identify specific DUBs involved in HCV replication by RNAi-based screening (Fig. 2A). The family of DUBs consists of approximately 100 genes. We established 61 stable Huh7.5.1 cell lines, each of which expressed an shRNA against one of the DUBs (Fig. 2A, Step 1) and selected 30 of the cell lines that exhibited an at least 40% reduction of DUBs expression (Fig. 2A, Step 2). We then inoculated HCV into the 30 cell lines and quantified the intracellular HCV RNA levels after 4 days (Fig. 2A, Step 3). We found that the cell lines harboring an shRNA against USP15 exhibited the most efficient reduction of HCV RNA in our screening (Fig. 2B). Next, we confirmed that USP15 expression was reduced in Huh7.5.1 cells expressing shRNA to USP15 (shUSP15) compared in Huh7.5.1 cells expressing control shRNA to LacZ (shLacZ) (Fig. 2C). In addition, the level of intracellular HCV RNA in Huh7.5.1 cells expressing shUSP15 upon infection with HCV was significantly lower than that in the Huh7.5.1 cells expressing shLacZ (Fig. 2D). These data suggest that USP15 is involved in the propagation of HCV.

**Figure 2.**
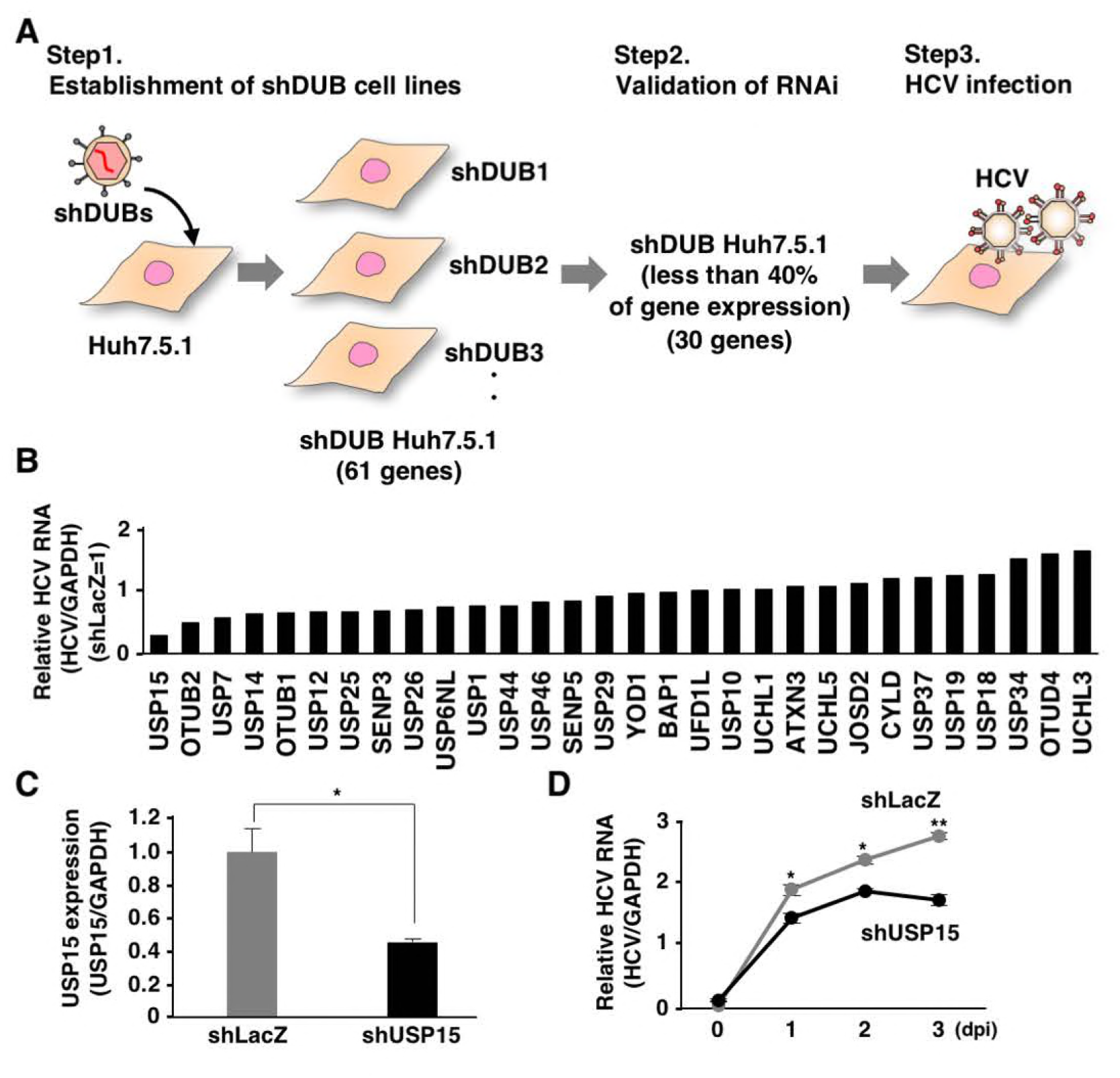
RNAi screening to identify specific DUBs involved in HCV propagation. (A) A schematic representation of the experimental procedure for our RNAi screening is shown. Huh7.5.1 cells were infected with retroviruses expressing shRNA targeting DUBs (shDUBs). The expression of the DUB gene was confirmed in each of the Huh7.5.1 cell lines expressing shDUBs. The shDUB Huh7.5.1 cell lines that exhibited a more than 40% reduction in the expression of their specific DUB were selected for further screening. DUB-knockdown Huh7.5.1 cells were infected with HCV at an moi of 0.5, and intracellular HCV RNAs were quantified after 4 days. (B) The levels of intracellular HCV RNAs at 4 days post-infection were determined by qPCR as a relative value against GAPDH mRNA in cells. Data are presented as the relative values compared to those in Huh7.5.1 cells expressing shRNA against the LacZ gene. (C) The expression of USP15 in Huh7.5.1 cells expressing shRNA targeting LacZ (shLacZ) or USP15 (shUSP15) was quantified by qPCR. (D) Huh7.5.1 cells expressing shRNA targeting LacZ or USP15 were infected with HCV at an moi of 0.5. The levels of intracellular HCV RNAs were determined by qPCR at the indicated time points.

### Deficiency of USP15 impairs HCV propagation

To further confirm the effect of USP15 on HCV propagation, we established two USP15-knockout Huh7 cell lines (USP15KOHuh7 #8 and #22; Fig. 3A) using the CRISPR/Cas9 system. The cell growth (Fig. 3B) and expression of miR-122, a determinant of HCV propagation in hepatocytes (Fig. 3C), of the USP15KOHuh7 cell lines were comparable to those of parental Huh7 cells. To evaluate the effect of USP15 deficiency on HCV propagation, Huh7 and USP15KOHuh7 cells were infected with HCV. Intracellular HCV RNA (Fig. 3D), an infectious titer in the culture supernatants (Fig. 3E) and the ratio of extra- and intracellular HCV RNA (Fig. 3F) at 4 days post-infection were decreased in USP15KOHuh7 cells. Exogenous expression of USP15 in USP15KOHuh7 cells restored the production of infectious HCV particles (Fig. 3G, H). These data suggest that USP15 is involved in HCV propagation. In addition, Huh7 and USP15KOHuh7 cells were transfected with an HCV subgenomic replicon RNA by electroporation, and colony formation was determined. Deficiency of USP15 decreased the numbers of colonies compared to those in Huh7 cells (Fig. 3I). To further examine the roles of USP15 in translation of HCV RNA, *in vitro* transcribed HCV subgenomic RNA possessing mutation in the active sites of RNA-dependent RNA polymerase of NS5B and a secreted form of NanoLuc (NLuc) as a reporter (pSGR-NLuc-JFH1GND) was transduced into Huh7 and USP15KOHuh7 cells by electroporation. NLuc activities were significantly suppressed in the USP15KOHuh7 cell lines (#8 and #22) compared with those in parental Huh7 cells (Fig. 3J). On the other hand, the activity of cap-dependent translation exhibited no significant difference between Huh7 and USP15KOHuh7 cells (Fig. 3K). Collectively, these results suggest that USP15 participates in HCV propagation through at least two distinct pathways, production of infectious particles and translation of viral RNA.

**Figure 3.**
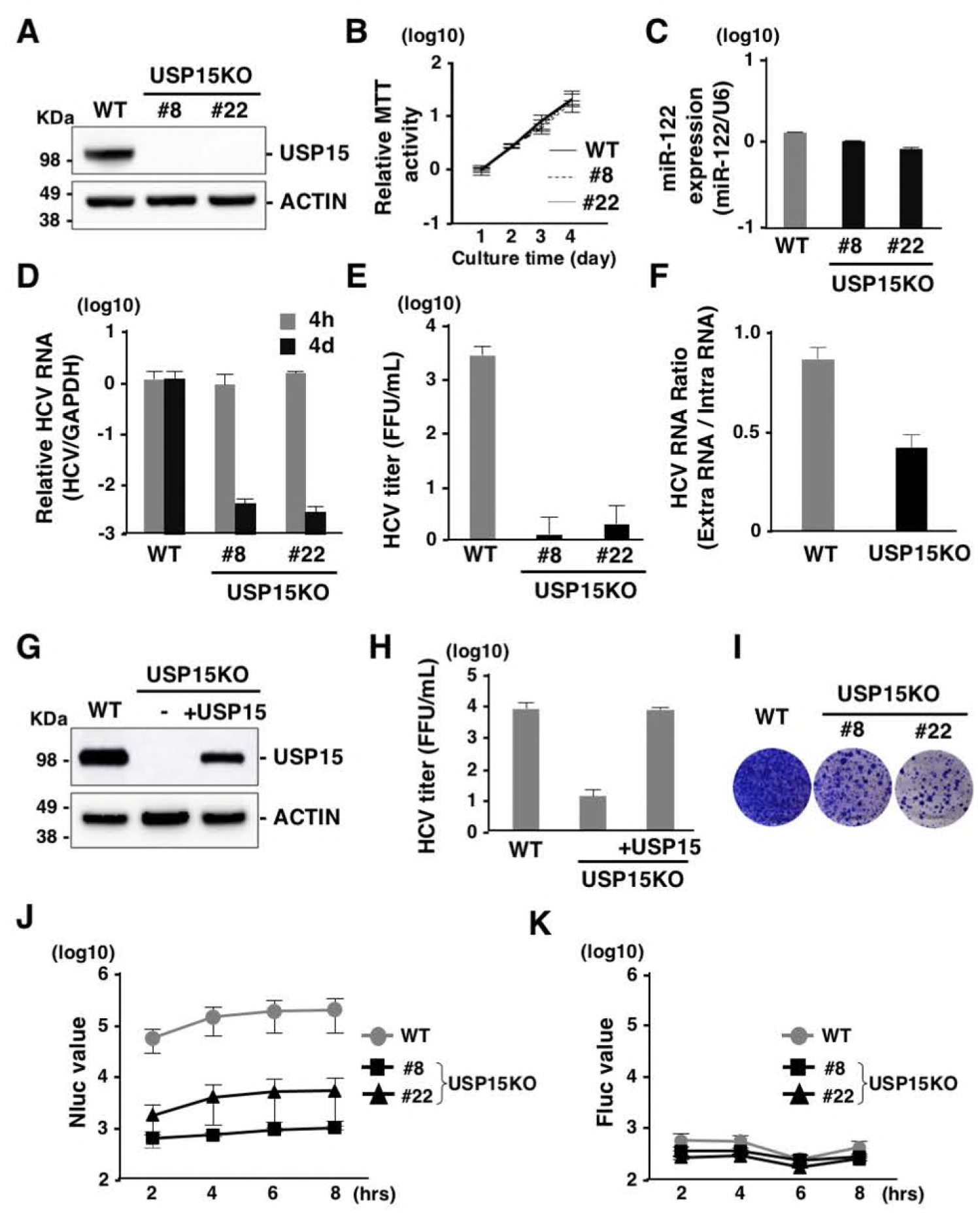
Deficiency of USP15 impaired HCV propagation. (A) USP15-deficient Huh7 (USP15KOHuh7) cells were generated using a CRISPR/Cas9 system. The expression of USP15 was confirmed by immunoblotting. We successfully established two independent USP15KOHuh7 clones. (B) WT and USP15KOHuh7 cells were seeded onto 24-well plates and incubated for 4 days. The cell growth of each cell line was analyzed by using an MTT assay kit (Nacalai Tesque, Kyoto, Japan) according to the manufacturer’s protocol. (C) The expression of miR-122 in WT and USP15KOHuh7 cells was quantified by qPCR. U6 RNA was used as the internal control. (D) HCV was infected into WT and USP15KOHuh7 cells at an moi of 3. After 4 h or 4 days post-infection, intracellular HCV RNA was quantified by qPCR using GAPDH mRNA as the internal control. (E) WT and USP15KOHuh7 cells were infected with HCV at an moi of 3. After 4 days post-infection, infectious titers in the culture supernatants were determined by focus-forming assay. (F) The WT and USP15KOHuh7 cells were infected with HCV at an moi of 3. After 4 days post-infection, extracellular and intracellular HCV RNAs were quantified by qPCR. The ratios between intracellular and extracellular HCV RNA were calculated as HCV RNA rates. (G) USP15KOHuh7 cells were transfected with a plasmid expressing FLAG-USP15, and USP15KOHuh7 cells expressing FLAG-USP15 cells were established. The expression of USP15 was confirmed by immunoblotting. (H) USP15KOHuh7, those with restored USP15 and Huh7 cells were infected with HCV. After 4 days post-infection, infectious titers in the culture supernatants were determined by focus-forming assay. (I) *In vitro* transcribed HCV subgenomic replicon RNA (pSGR-JFH1) was electroporated into WT and USP15KOHuh7 cells, and the cells were incubated for 3 weeks in the presence of 1 mg/mL of G418. Colonies were visualized by Giemsa staining. (J) *In vitro* transcribed HCV subgenomic replicon RNA (pSGR-NLuc-JFH1GND) was electroporated into WT and USP15KOHuh7 cells, and the activity of Nluc in culture supernatants was monitored. (K) *In vitro* transcribed capped-Fluc RNA was electroporated into parental and USP15KO Huh7 cells and the activity of Fluc in electroporated cells was monitored.

### USP15 supports HCV propagation in a hepatocyte-specific manner

Not only Huh7 cells but also Hep3B cells expressing miR-122 (Hep3B/miR-122), HepG2 cells expressing CD81 (HepG2/CD81) and 293T cells expressing miR-122, Claudin 1 (CLDN1) and APOE (293T/miR-122/CLDN1/APOE) have been shown to permit HCV propagation (36–38). To examine the effects of USP15 on HCV propagation in these cell lines, we established USP15KO cell lines in Hep3B/miR122 and 293T/miR-122/CLDN1/APOE cells (Fig. 4A and 4B). We could not obtain USP15-knockout HepG2/CD81 cells (data not shown). USP15KO and parental Hep3B/miR-122 cells were infected with HCV and intracellular HCV RNA was determined at 4 h and 4 days post-infection. Although intracellular HCV RNA levels were comparable in USP15KO and parental Hep3B/miR-122 cells at 4 h post-infection, they were significantly reduced in USP15KO cells at 4 days post-infection (Fig. 4C). Infectious titers in the culture supernatant at 4 days post-infection were also reduced in USP15KO cells (Fig. 4E). In contrast, intracellular HCV RNA in USP15KO and parental 293T/miR-122/CLDN1/APOE cells at 12 h and 2 days post-infection (Fig. 4D) and infectious titers in the culture supernatant at 2 days post-infection (Fig. 4F) were comparable. To examine the effect of USP15 in 293T cells on translation of viral RNA, subgenomic HCV RNA of SGR-Nluc-JFH1GND was electroporated into USP15KO and parental 293T cells expressing miR-122. Although the expression of miR-122 in parental 293T cells significantly enhanced the translation efficiency of HCV RNA, Nluc activities were comparable among USP15KO and parental 293T cells expressing miR-122 (Fig. 4G), suggesting that USP15 participates in HCV propagation in an miR-122-independent and hepatocyte-specific manner.

**Figure 4.**
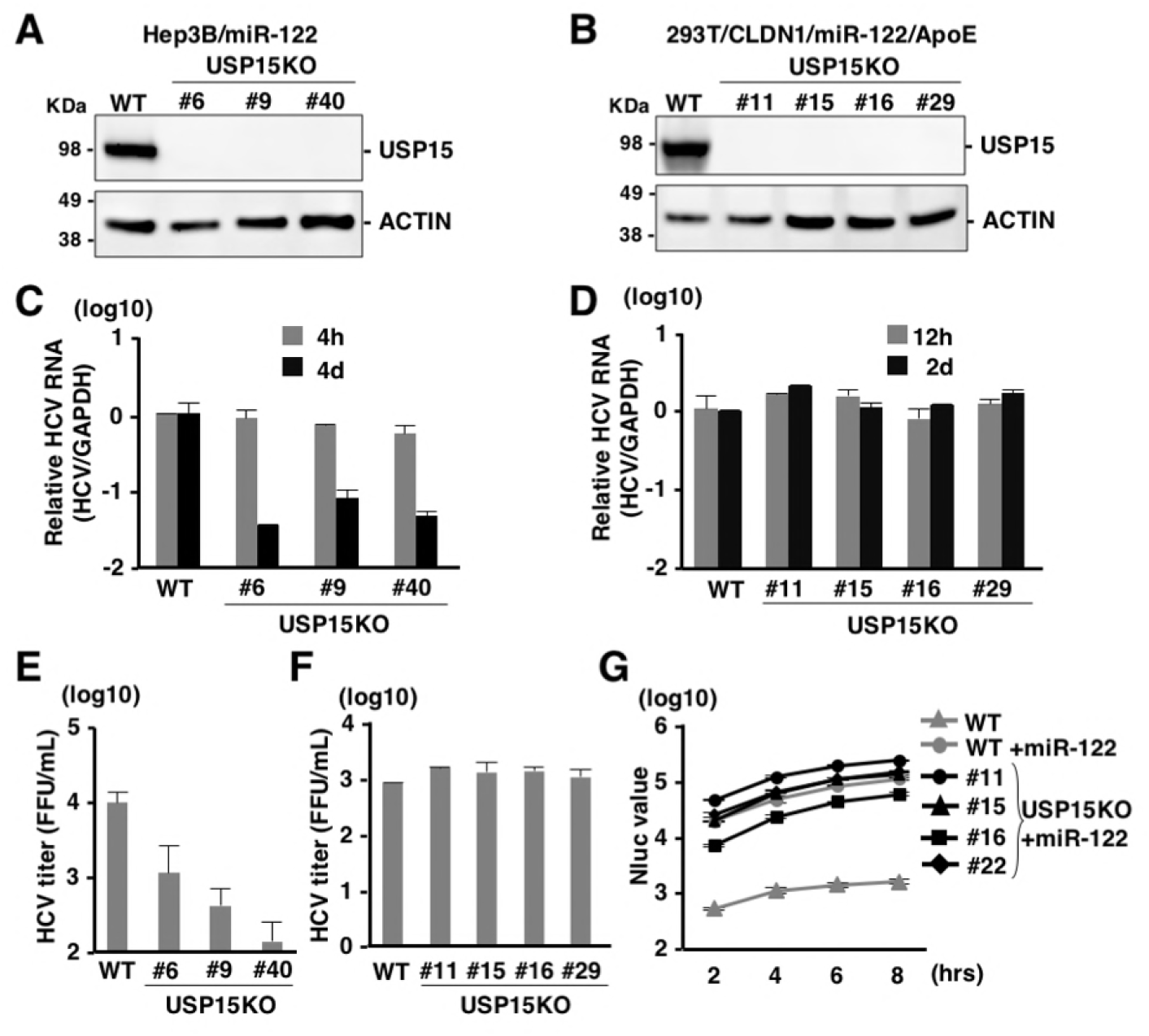
Cell-type-specific reduction of HCV propagation in USP15KO cells. (A) USP15KOHep3B/miR122 cells were generated using the CRISPR/Cas9 system. The expression of USP15 was confirmed by immunoblotting. (B) USP15KO293T cells were also generated using the CRISPR/Cas9 system, and subjected to immunoblotting to confirm the USP15 expression. (C) USP15KOHep3B/miR-122 and parental Hep3B/miR-122 cells were infected with HCV at an moi of 3. After 4 h or 4 days post-infection, intracellular HCV RNA was quantified by qPCR using GAPDH mRNA as the internal control. (D) Parental and USP15KO 293T cells were lentivirally transduced with miR-122, CLDN1 and APOE, and then infected with HCV (moi=10) at 2 days post-transduction. After 12 h or 2 days post-infection, intracellular HCV RNAs were quantified by qPCR. (E) Infectious titers of parental and USP15KO Hep3B/miR-122 cells in the culture supernatants were determined by focus forming assay at 4 days post-infection. (F) Infectious titers of parental and USP15KO 293T cells in the culture supernatants were determined by focus forming assay at 2 days post-infection. (G) *In vitro* transcribed subgenomic HCV replicon RNA (SGR-NLuc-JFH1GND) was electroporated into parental and USP15KO 293T cells expressing miR-122, and the activity of Nluc in the culture supernatants was determined.

### USP15 is not involved in innate immune responses

USP15 has been reported as a DUB targeting tripartite motif-containing protein 25 (TRIM25) and positively regulated RIG-I-mediated innate immune responses (39). TRIM25 has been shown to conjugate the K63-linked ubiquitin chains to RIG-I to facilitate downstream signaling pathways (39). USP15 has reported to remove the K48-linked ubiquitin chains of TRIM25 mediated by the linear ubiquitin assembly complex (LUBAC) to protect TRIM25 from proteasomal degradation (39). On the other hand, USP15 has been shown to target RIG-I and to remove the K63-linked ubiquitin chains from RIG-I (40). RIG-I senses viral RNAs such as Japanese encephalitis virus (JEV) and vascular stomatitis virus (VSV) and activates downstream molecules to activate innate immune responses (41). In contrast, encephalomyocarditis virus (EMCV) is recognized by melanoma differentiation-associated gene 5 (MDA5) rather than RIG-I (41). To investigate the involvement of USP15 in RNA virus infection, parental and USP15KOHuh7 cells were infected with JEV, VSV and EMCV. Although intracellular JEV RNA levels were comparable between parental and USP15KO cells (Fig. 5A), infectious titers in the culture supernatants at 2 days post-infection were slightly decreased in USP15KO cells (Fig. 5B). In contrast, viral titers of VSV and EMCV were comparable between parental and USP15KO cells (Fig. 5C and 5D). These data suggest that USP15 plays a small role in the propagation of JEV but is not involved in the propagation of VSV and EMCV.

**Figure 5.**
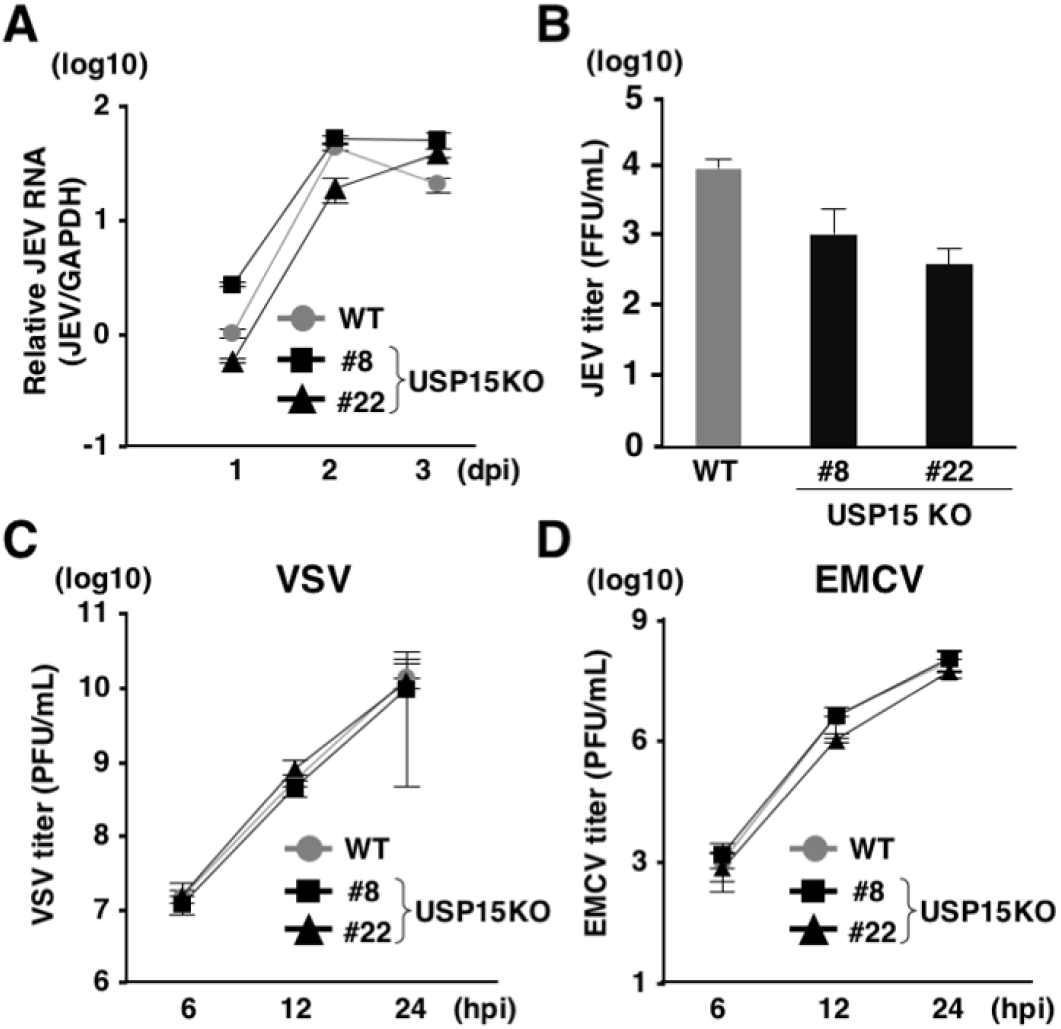
USP15 is partially involved in propagation of JEV but not VSV or EMCV. (A) WT or USP15KOHuh7 cells were infected with JEV at an moi of 3. Intracellular JEV RNA was quantified by qPCR at each time point. (B) WT or USP15KOHuh7 cells were injected with JEV at an moi of 3 and incubated for 2 days. Infectious JEV titers in the culture supernatants were determined by focus forming assay. (C) WT and USP15KOHuh7 cells were infected with VSV at an moi of 3. Infectious VSV titers in the culture supernatants were determined by plaque forming assay at the indicated time points. (D) WT and USP15KOHuh7 cells were infected with EMCV at an moi of 1. Infectious EMCV titers in the culture supernatants were determined by plaque forming assay at the indicated time points.

To further assess the involvement of USP15 in RIG-I-mediated antiviral effects *in vivo*, we generated USP15^-/-^ mice possessing deletion of 223 base pairs in the *Usp15* genomic locus by using the CRISPR/Cas9 system (Fig. 6A). The USP15^-/-^ mice were fertile and visually normal as reported previously (42). We intranasally challenged the USP15^-/-^, USP15^+/+^ and IFNα/βR^-/-^ mice with a lethal dose of VSV and monitored the survival rates and body weights. Deficiency of USP15 had no significant effect on the survival of mice against VSV infection, while IFNα/βR^-/-^ mice showed high sensitivity to VSV challenge (Fig. 6B) and the change of body weight was comparable between USP15^+/+^ and USP15^-/-^ mice (Fig. 6C), suggesting that USP15 does not participate in survival after VSV challenge. In addition, mouse embryonic fibroblasts (MEFs) prepared from USP15^+/+^ and USP15^-/-^ mice were infected with VSV and induction of ISGs was monitored. The inductions of *Ifnα*, *Cxcl10* and *Il6* were comparable between the two MEFs (Fig. 6E to 6G). These results suggest that innate immune responses do not participate in the USP15-mediated enhancement of HCV propagation.

**Figure 6.**
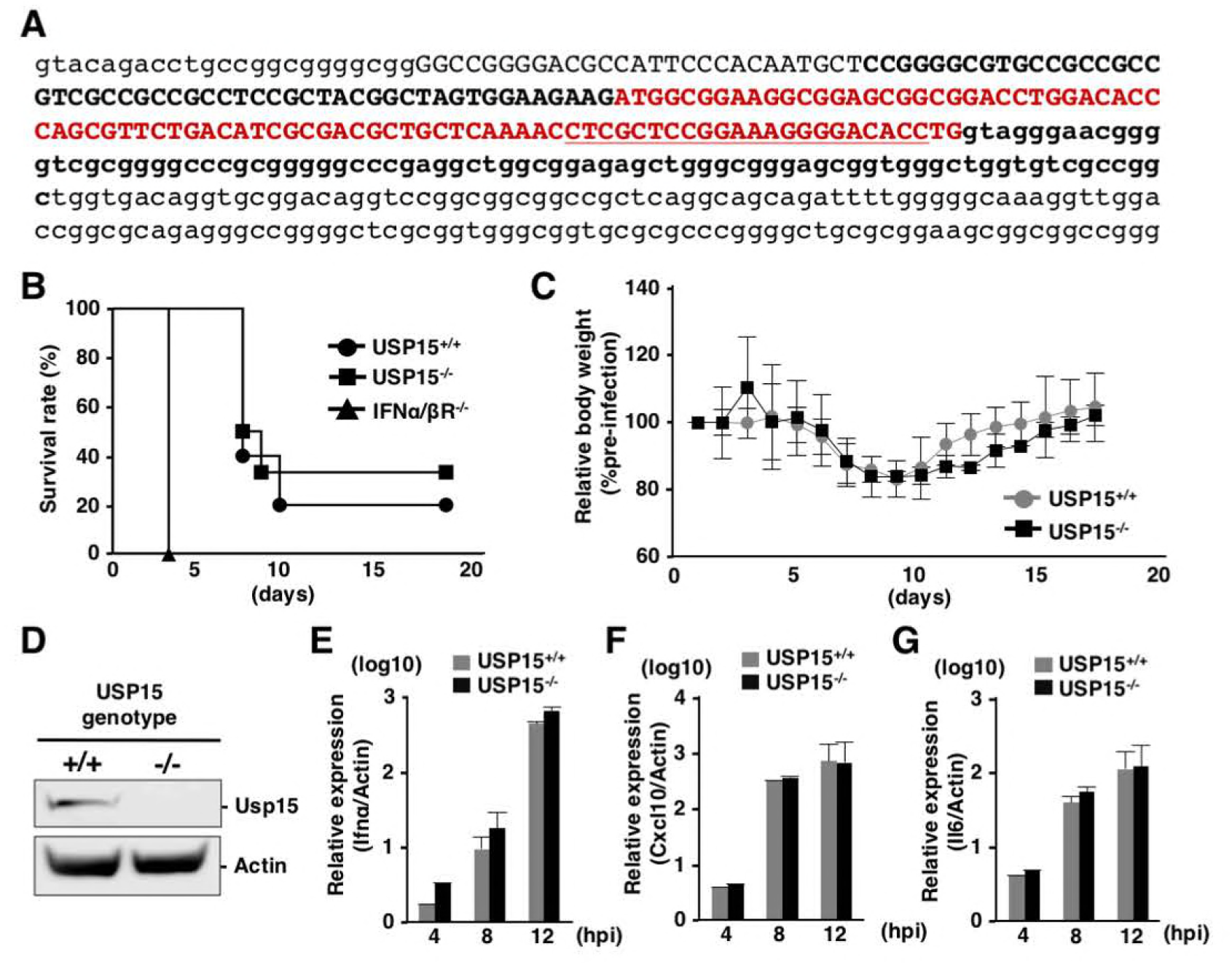
USP15 is not involved in innate immune responses *in vivo*. (A) Generation of USP15^-/-^ mice. Small letters indicate parts of introns and capital letters indicate exon 1. The red color sequence is the open reading frame of exon 1 of USP15. Sequences of the guide RNA targeting USP15 are underlined. USP15^-/-^ mice possessed the 223-nucleotide deletion shown in bold. (B, C) USP15^+/+^ (N=10), USP15^-/-^(N=6) and IFNα/ßR^-/-^ (N=5) mice (16–17 weeks old) were intranasally infected with VSV (4 × 10^6^ pfu) and their survival rates (B) and body weights (C) were monitored daily. Body weight changes at each time point were indicated as the relative values against the body weights of mice at 1 day post-infection. (D) MEFs were prepared from USP15^-/-^ and USP15^+/+^ mice. The expression of USP15 was confirmed by immunoblotting. (E, F, G) MEFs infected with VSV at an moi of 1 were collected at each time point and the mRNAs of *Ifnα* (E), *Cxcl10* (F) and *Il6* (G) were quantified by qPCR.

### USP15 participates in lipid metabolism to facilitate HCV propagation

Due to a lack of information about the function of USP15 in hepatocytes, we investigated the roles of USP15 during HCV infection. First, we investigated the possibility that USP15 interacts with viral proteins. Immunoprecipitation analysis revealed no interaction of USP15 with viral proteins (Fig. 7A). Confocal microscopic observation showed that USP15 co-localized with LDs (Fig. 7B). Previous reports suggest that LDs and lipid metabolism are important for HCV replication and infectious particle formation (19, 24, 43). Therefore, we investigated the roles of USP15 on formation of LDs. LDs in parental and USP15KOHuh7 cells were stained with HCS LipidTOX^TM^ Red neutral lipid stain and observed by confocal microscopy. The amounts and sizes of LDs were significantly reduced in USP15KO cells compared with parental Huh7 cells (Fig. 7C and 7D). The adipose differentiation-related protein (ADRP) is a marker for LDs (44), and expression of ADRP was significantly impaired in the USP15KOHuh7 cell lines (Fig. 7E). LDs consist of neutral lipids such as cholesterol esters and triglycerides (TGs) and function as storage sites for fatty acids (45). SREBP-1c and SREBP-2 are master transcriptional factors that regulate fatty acid synthesis and cholesterol synthesis, respectively (46, 47). Next, we examined the expression of these transcriptional factors in USP15KOHuh7 cells. Expression of SREBP-1c but not SREBP-2 was significantly reduced in USP15KOHuh7 cells (Fig. 7F and 7G), suggesting that reduction of fatty acids production participates in the suppression of HCV propagation in USP15KOHuh7 cells. To verify this possibility, palmitic acid (PA) and oleic acid (OA) were added to the culture media in parental and USP15KOHuh7 cells, and intracellular HCV RNA was determined at 4 days post-infection with HCV. Although intracellular HCV RNA levels in both parental and USP15KOHuh7 cells exhibited no significant change by the addition of fatty acids (Fig. 7H and 7I), infectious titers in the culture supernatants of USP15KO but not those of parental Huh7 cells were significantly enhanced by the addition of PA, but not of OA (Fig. 7J and 7K). Taken together, these results suggest that USP15 participates in the regulation of lipid metabolism and facilitates production of infectious HCV particles.

**Figure 7.**
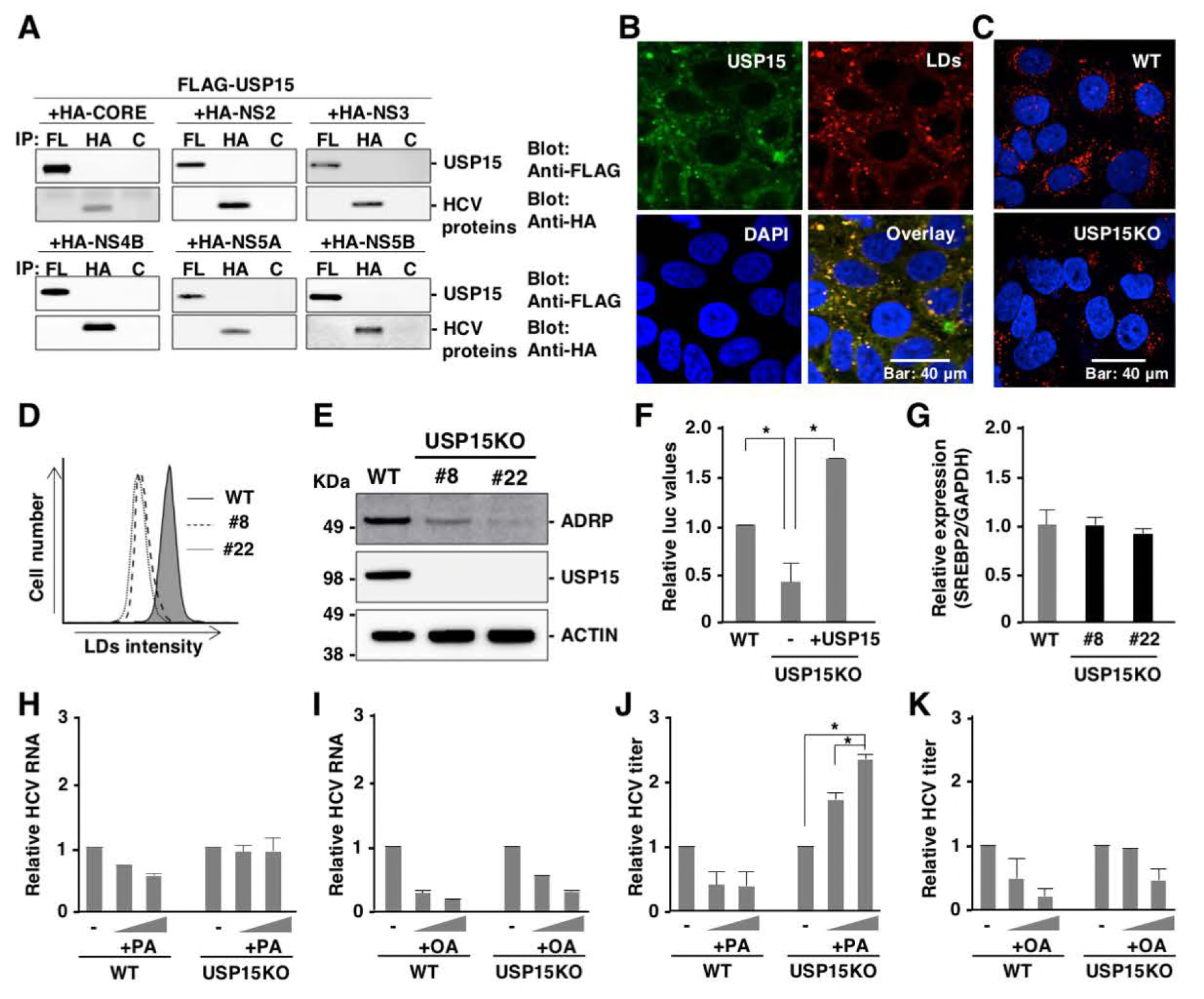
USP15 controls LD formation to facilitate HCV propagation. (A) Interactions between FLAG-USP15 and HA-tagged viral proteins in 293T cells were evaluated by immunoprecipitation with antibodies to FLAG (FL)-, HA- and Glu/Glu-tag (Control, C). Immunoprecipitants were subjected to immunoblotting by using FLAG and HA antibodies. (B) Subcellular localization of USP15 in Huh7 cells was observed by confocal microscopy. USP15 (green), lipid droplets (red) or nuclei (blue) were stained with anti-USP15 antibody, HCS LipidTOX^TM^ Red neutral lipid stains and DAPI, respectively. (C) Lipid droplets in WT or USP15KOHuh7 cells were observed by confocal microscopy. Lipid droplets (red) or nuclei (blue) were stained with HCS LipidTOX^TM^ Red neutral lipid stains and DAPI, respectively. (D) The intensity of stained LDs was quantified by FACS. (E) The expression of ADRP, a marker of LDs, in parental and USP15KO Huh7 cells was detected by immunoblotting by using the indicated antibodies. (F) The expressions of SREBP-1c in the WT cells, USP15KO cells and restored USP15 Huh7 cells were examined by using reporter assay. Huh7 cells, USP15KOHuh7 and USP15KOHuh7 cells expressing USP15 were transfected with pGL3-Basic SREBP-1c and pRL-TK and incubated for 2 days. Luciferase activity was determined by using a Dual-Luciferase Reporter Assay System (Promega, Madison, WI). (G) Expression of SREBP2 in parental and USP15KO Huh7 cells was quantified by qPCR. mRNA of GAPDH was used as the internal control. (H, J) WT and USP15KO Huh7 cells were treated with 100 or 200 µM palmitic acid (PA) during HCV infection at an moi of 3. After 4 days post-infection, intracellular HCV RNA (H) or infectious HCV titers (J) in culture supernatants were determined. (I, K) WT and USP15KOHuh7 cells were treated with 100 or 200 µM oleic acid (OA) during HCV infection at an moi of 3. After 4 days post-infection, intracellular HCV RNA (I) or infectious HCV titers (K) in the culture supernatants were determined.

## Discussion

In this study, we identified USP15 as a novel host factor for HCV propagation. This is the first report to identify a specific DUB involved in HCV propagation. Our data suggest that USP15 participates in at least two steps in the HCV life cycle: translation of viral RNA and production of infectious particles. HCV RNA is translated to viral proteins through the internal ribosomal entry site (IRES) (48). Deficiency of USP15 significantly impaired the translation of viral RNA specific to HCV-IRES. In addition, lack of USP15 showed no effect on HCV propagation in non-hepatic cells such as 293T cells, suggesting that USP15 targets hepatocyte-specific factors to regulate translation of HCV RNA. Liver-specific miR-122 binds to the two sites of HCV RNA to facilitate HCV replication (11, 49) and many reports suggest that binding of miR-122 to HCV RNA promotes translation of HCV RNA (50, 51). Because DUB is unlikely to directly interact with miR-122, USP15 may interact with components of the miR-122/AGO2 complex. Although activity of HCV IRES-mediated RNA translation was significantly enhanced by the expression of miR-122 as previously reported, deficiency of USP15 did not affect the enhancement of HCV IRES-mediated translation by the expression of miR-122 (Fig 4G), suggesting that USP15 participates in HCV IRES-mediated translation through an miR-122 independent pathway. Various host factors, such as heterogeneous nuclear ribonucleoprotein L (hnRNP L), nuclear factor (NF) 90, NF45, poly-C binding protein 2 (PCBP2), and insulin-like growth factor 2 mRNA-binding protein 1 (IGF2BP1), have been shown to participate in IRES-mediated translation and replication of HCV through the interaction between 5’- and 3’-UTR of HCV (52–55). Further studies are needed to understand the roles of USP15 in HCV RNA translation through the interaction with these host factors.

LDs are cellular organelles for the storage of TGs and have a single membrane (56). Once cells require energy, stored TGs undergo hydrolysis to produce fatty acids through the activation of lipolytic pathways. A number of LD-associated proteins have been identified. The perilipin family—which includes perilipin, ADRP, tail-interacting protein of 47 kilodaltons (TIP47), and S3–12 and myocardial lipid droplet protein (MLDP)/oxidative tissues-enriched PAT protein (OXPAT)/lipid storage droplet protein 5 (LSDP5)—are major regulators of LD homeostasis (57, 58). Among them, HCV NS5A interacts with TIP47 to facilitate HCV replication (59). In addition, HCV core protein has been shown to displace ADRP from the LD surface (60) and to contribute to efficient virus assembly through interaction with LDs (61). In this study, we showed that USP15 co-localizes with LDs in Huh7 cells without any interaction with HCV proteins, suggesting that USP15 interacts with LD-associated proteins to regulate LD formation and is involved in viral assembly. We showed that lack of USP15 reduces the amounts of LDs in Huh7 cells. It is known that non-hepatic cells such as 293T cells possess small amounts of LDs, and in 293T cells LDs and LD-related fatty acids are not involved in the propagation of HCV. This might be one of the reasons that HCV propagation in USP15KO293T cells is comparable to that in parental cells. We also showed that expression of SREBP-1c was impaired in USP15KOHuh7 cells. It was demonstrated that HCV core protein enhances the binding of liver X receptor α 264 (LXRα) and retinoid X receptor α (RXRα) to LXR-response element during HCV infection (62, 63). These reports indicate that HCV core protein plays crucial roles in the modulation of lipid metabolism during HCV infection. However, immunoprecipitation analyses showed that USP15 does not interact with HCV core protein, suggesting that USP15 regulates lipid metabolism independent of the interaction with HCV core protein.

USP15 is expressed in various tissues and has been shown to be involved in various cellular events. USP15 targets receptor-activated SMADs (R-SMADs) to regulate TGF-β signaling (64). In addition, USP15 interacts with SMAD7 and SMAD-specific E3 ubiquitin protein ligase 2 (SMURF2) and de-ubiquitinates type I TGF-β receptor (65). USP15 has been reported to de-ubiquitinate Kelch-like ECH-associated protein 1 (Keap1), which regulates NF-E2-related factor (Nrf) 2-dependent antioxidant responses (66), murine double minute 2 (MDM2) (42), histone H2B (H2B) (67), Nrf1 (68) and p62 (69). USP15 is also suggested to cleave ubiquitin chains of viral proteins such as Nef of HIV-1 (70), HBx of HBV (71) and E6 protein of human papillomavirus (72). These reports indicate that USP15 participates in the development of many types of cancers, cellular homeostasis and virus infection. However, the substrates and function of USP15 in hepatocytes have not been clarified. We showed that the birth of USP15^-/-^ mice followed Mendelian ratios, and the mouse pups developed normally and exhibited no obvious abnormalities, as reported previously (42). The phenotypes of USP15^-/-^ mice and our data suggest that there are unknown substrates of USP15 which regulate lipid metabolism. Recently, USP15 has been shown to be a DUB for TRIM25 (39) and RIG-I (40) and to be involved in type I IFN response in cells infected with RNA viruses. Upon infection of HCV, viral RNA is sensed by RIG-I and type I IFN and ISG are induced (73). We examined the involvement of USP15 in innate immune responses and propagation of various RNA viruses and revealed that USP15 does not participate in either the survival of mice or innate immune responses in MEFs. These data suggest that USP15 is dispensable for the induction of innate immune responses upon infection with RNA viruses. Further characterization of USP15^-/-^ mice is needed in order to elucidate the physiological function of USP15, especially in hepatocytes.

Fatty acids have been shown to support HCV replication in replicon cells (74, 75). On the other hand, several recent papers showed that fatty acids inhibit HCV replication (74, 76–78). In the present study, we found that deficiency of USP15 in Huh7 cells reduced the size and number of LDs, and addition of PA—but not of OA—partially restored the production of infectious HCV particles. We do not have any data indicating why OA did not support the production of infectious HCV. However, our data do suggest that PA and its metabolites support the production of infectious HCV particles. On the other hand, we could not observe the enhancement of HCV particle production by the treatment with PA in naive Huh7 cells due to the abundance of LDs. Taken together, our data suggest that USP15 mainly contributes to replication of HCV RNA and partially supports HCV assembly.

In summary, we identified USP15 as a novel host protein involved in HCV propagation. USP15 participates in HCV translation and plays a role in the production of infectious particles. Our data suggest that USP15 participates in the formation of LDs through the regulation of hepatic lipid metabolism to facilitate HCV propagation. Because USP15 possesses the enzymatic activity of de-ubiquitinatinase, and thus removes ubiquitin from substrates, in a future study it would be of interest to identify the substrates of USP15 in hepatocytes. The hepatocyte-specific substrates of USP15 that are crucial for HCV propagation might be novel drug targets for chronic hepatitis C.

## Materials and Methods

### Cell lines and viruses

Huh7, Huh7.5.1, Hep3B/miR122, Vero, 293T, Plat-E and BHK-21 cells were obtained from the National Institute of Infectious Diseases and were cultured in Dulbecco’s Modified Eagle’s medium (DMEM) supplemented with 10% fetal bovine serum (FBS), 100 U/ml penicillin and 10 µg/ml streptomycin. HCV replicon cells (9–13) (79) were maintained in DMEM supplemented with 10% FBS, 1 mg/ml G418, 100 U/ml penicillin and 10 µg/ml streptomycin. HCV derived from the genotype 2a JFH-1 strain mutated in E2, p7 and NS2 was prepared by serial passages in Huh7.5.1 cells as mentioned previously (80). JEV (AT31 strain) was propagated in C6/36 cells. VSV and EMCV were propagated in BHK-21 cells.

### Antibodies and reagents

The following antibodies were used: anti-JEV NS3 monoclonal antibody (#578) (81), anti-HCV NS5A monoclonal antibody (5A27) (82), anti-ACTIN mouse monoclonal antibody (A2228, Sigma-Aldrich), horseradish peroxidase-conjugated anti-FLAG mouse monoclonal antibody (A8592, Sigma-Aldrich), anti-USP15 monoclonal antibody (ab56900, abcam), anti-FLAG mouse monoclonal antibody (F1804, Sigma-Aldrich), anti-HA monoclonal antibody (clone 16B12, MMS-101P, Biolegend) and anti-Glu-Glu antibody (MMS-115P, Biolegend). Oleic acid (O1008) and Palmitic acid (P0500) were obtained from Sigma-Aldrich. PR-619 (SI9619) was purchased from Lifesensor. HCS LipidTOX^TM^ Red neutral lipid stain was obtained from Thermo Fisher Scientific.

### Plasmids

The shRNAs against each DUB were obtained from the Human shRNA library (Takara Bio). The pRSV-Rev (#12253), pMDLg/pRRE (#12251), pCMV-VSV-G (#8454), pX330 (#42230), and pCAG EGxxFP (#50716) were obtained from Addgene. Reporter plasmid, pGL3-Basic SREBP1c and pRL-TK were previously used (62). Lentiviral vectors expressing miR-122, APOE and Claudin 1 (CLDN1) were also used as described previously (19). The cDNAs of USP15, OTUB1, OTUD1 and OTUD7B obtained from Dr. Wade Harper (Addgene; #22570, #22551, #22553 and #22550) were amplified by PCR and cloned into pEF FLAG pGK puro (83) or FUIPW (26) by using an In-Fusion HD cloning kit (Takara Bio). HCV core, NS2, NS3, NS4B, NS5A and NS5B were amplified by PCR and cloned into pCAGGS (84). The sgRNAs of human USP15 (5’-CACCGCGACTATCGACTAGGTACC-3’) and mouse USP15 (5’-CACCGGTGTCCCCTTTCCGGAGCG-3’) were cloned into pX330 by Ligation high Ver. 2 (Toyobo). Genomic DNAs of Huh7 cells or mouse tails were extracted by DirectPCR Lysis Reagents (Viagen Biotech), and amplified by PCR using the following primers sets: human USP15 (forward, 5’-CAACCACTGAGGATCCGCTCCCGGTGTCTTTTGGTTTCGA-3’; reverse, 5’-TGCCGATATCGAATTCCCTATCATTCGGGAAGGCCTGAGGT-3’) and mouse USP15 (forward, 5’-CAACCACTGAGGATCCATTTGGTACAGACCTGCCGG-3’; reverse, 5’-TGCCGATATCGAATTCTCGGAATAATGGGGAACTTGGG-3’). They were then cloned into pCAG EGxxFP by using an In-Fusion HD cloning kit. The pSGR-JFH1 was mutated in the GDD motif of NS5B to GND to generate an inactive form of RNA-dependent RNA polymerase. In addition, the cDNA of NanoLuc was replaced with the neo gene to generate pSGR-NLuc-JFH1GND. The cDNA of firefly luciferase (FLuc) was amplified and cloned into pCMVTNT vector (Promega) designed as pCMVTNT Fluc. All plasmids used in this study were confirmed by sequencing with an ABI Prism 3130 genetic analyzer (Thermo Fisher Scientific).

### RNAi screening

Retroviruses expressing shRNAs against human DUBs were generated in Plat-E cells. Briefly, Plat-E cells (2 × 10^6^ cells) were seeded on a 10 cm dish and incubated at 37℃ for 24 h. 5 µg of retroviral transfer vector and 1 µg of pCMV-VSV-G were mixed with 500 μl of Opti-MEM and 40 μl of polyethylenimine (1 mg/ml; Polysciences) and incubated for 15 min. The DNA complex was inoculated into Plat-E cells, and the culture medium was changed at 4 h post-transfection. The culture supernatants were collected at 3 days post-transfection. Huh7.5.1 cells (2 × 10^5^ cells per well) were seeded on six-well plates and were incubated for 24 h. The virus-containing culture supernatants (2 ml) and 8 μl of Polybrene (4 mg/ml; Sigma-Aldrich) were inoculated into Huh7.5.1 cells and centrifuged at 1,220 × g for 45 min at 32 °C. Stable cell lines were selected by puromycin at 2 days post-infection. To determine the efficiency of RNAi, the expression of each DUB was analyzed by using real-time PCR (qPCR) as described below. Huh7.5.1 cells expressing shRNA were seeded on 24-well plates (3 × 10^4^ cells per well), incubated for 24 h and inoculated with HCV at a multiplicity of infection (moi) of 0.5. After 2 h, the culture medium was changed and the infected cells were incubated for 4 days. Intracellular HCV RNA was quantified by qPCR.

### qRT-PCR

Total RNA was extracted by using ISOGEN II (Nippon Gene). Real-time PCR for HCV or JEV RNA was performed by using a TaqMan RNA-to-Ct 1-Step Kit and a ViiA7 real-time PCR system (ThermoFisher Scientific). The following primers were used: HCV, 5′-GAGTGTCGTGCAGCCTCCA-3′ and 5′-CACTCGCAAGCACCCTATCA-3′; JEV, 5′-GGGTCAAAGTCATTTCTGGTCCATA-3’ and 5′-TCCACGCTGCTCGAA-3’; GAPDH, 5′-TGTAGTTGAGGTCAATGAAGGG-3′ and 5′-ACATCGCTCAGACACCATG-3′. The following probes were used: HCV, 5′-6-FAM/ CTGCGGAAC/ZEN/CGGTGAGTACAC/-3′IABkFQ; JEV, 5’-6-FAM/ATGACCTCG/ZEN/CTCTCCC/-3’IABkFQ; GAPDH, 5′-6-FAM/AAGGTCGG-A/ZEN/GTCAACGGATTTGGTC/-3′IABkFQ. Relative amounts of HCV or JEV RNA were determined by the ΔΔCt method using GAPDH as an internal control. For gene expression analysis, qPCR was performed by using a Power SYBR Green RNA-to Ct 1-step Kit (Thermo Fisher Scientific). The primers used in this study are summarized in Table 1.

**Table. 1.**
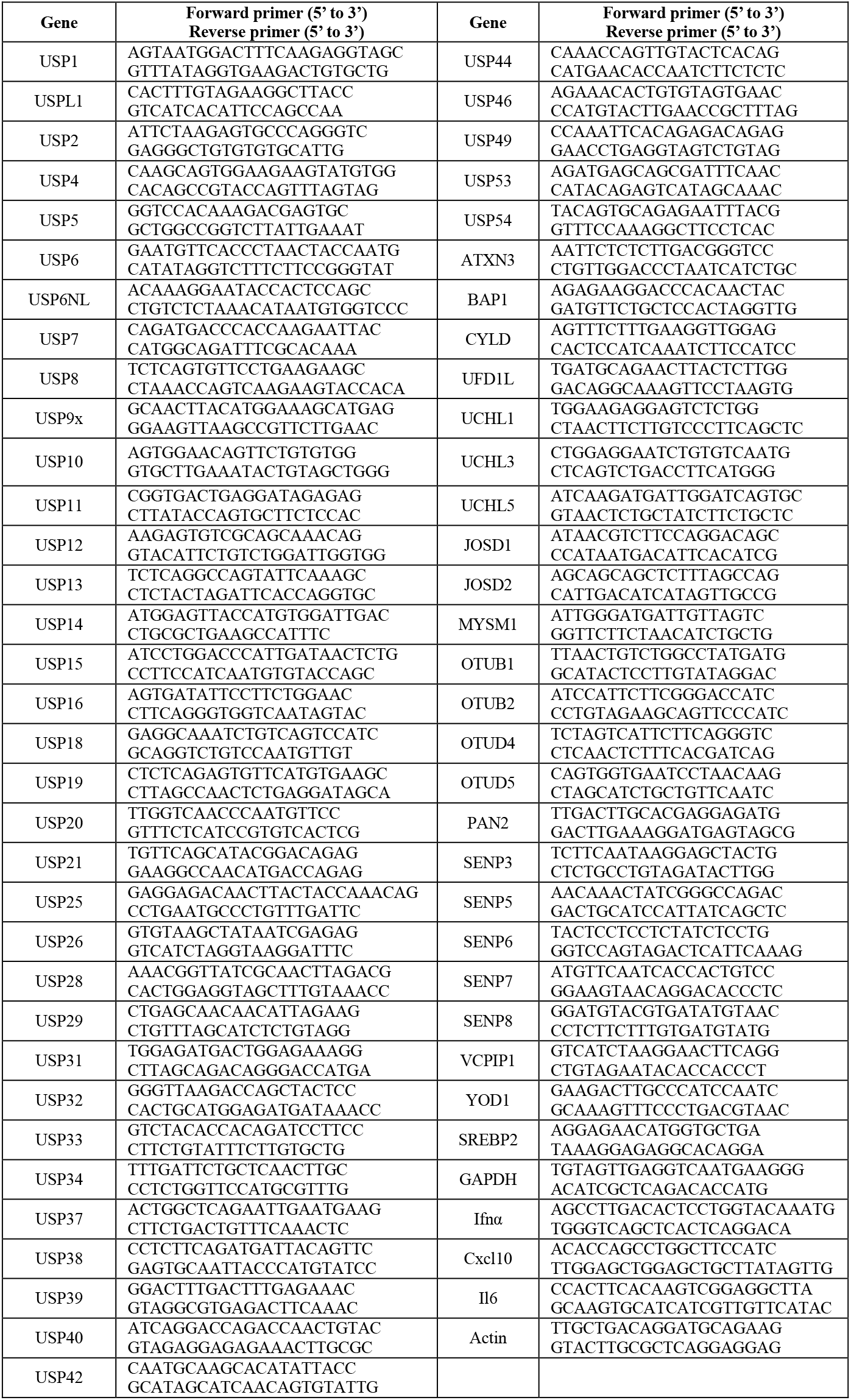
Primer set list for real-time PCR.

### Cell viability assay

Huh7 (9–13) cells were seeded on 24-well plates (5 × 10^4^ cells per well) and incubated at 37℃ for 24 h. PR-619 (0.5 µM) was added and the cells were incubated for an additional 24 h. Supernatants containing cells and adherent cells were collected and stained with 5 μg/ml of propidium iodide (PI, P4170; Sigma-Aldrich). Cell viability was determined by flow cytometry analyses (BD) using FlowJo software (FlowJo).

### Generation of USP15-knockout Huh7, Hep3B and 293T cells

Gene-knockout Huh7, Hep3B/miR122 and 293T cells were generated by using the CRISPR/Cas9 system as previously described (85). Briefly, cells were transfected with pX330 and pCAG EGxxFP and incubated for 1 week. GFP-positive cells were sorted by FACS and formed single colonies. Gene deficiency was confirmed by sequencing and western blotting.

### *In vitro* transcription, RNA transfection and colony formation

The plasmid pSGR-JFH1 was linearized with XbaI and transcribed *in vitro* by using a MEGAscript T7 kit (ThermoFisher Scientific) according to the manufacturer’s protocol. The pCMVTNT Fluc was linearized with BamHI and transcribed *in vitro* by using an mMESSAGE mMACHINE T7 ULTRA Transcription Kit (ThermoFisher Scientific) according to the manufacturer’s protocol. The *in vitro* transcribed RNA (10 μg) was electroporated into Huh7 cells (5 × 10^6^ cells) under conditions of 270 V and 950 μF using a Gene Pulser apparatus (Bio-Rad). For the transient experiments, cells were added into 10 ml of culture medium and plated on 12-well plates. For long-term colony formation, electroporated cells were plated in DMEM containing 10% FBS. The medium was replaced with fresh DMEM containing 10% FBS and 1 mg/ml G418 at 24 h post-electroporation. Colonies were visualized by staining with Giemsa (Merck) at 3 weeks post-electroporation.

### Immunofluorescence staining

Huh7 cells were fixed with 4% paraformaldehyde in phosphate-buffered saline (PBS) for 2 h. Cells were washed by PBS and permeabilized by 0.2% TritonX-100 in PBS for 15 min. After washing with PBS, the fixed cells were incubated with anti-NS5A mouse monoclonal antibody or anti-USP15 mouse monoclonal antibody at room temperature for 1 h. After washing, cells were incubated with Alexa Fluor (AF) 488-conjugated anti-mouse antibody and HCS LipidTOX^TM^ Red neutral lipid stain diluted by 2% FBS in PBS at room temperature for 1 h. The stained cells were covered with Prolong Gold AntiFade Reagent with DAPI (Thermo Fisher Scientific) and observed by FluoView FV1000 confocal microscopy (Olympus).

### Immunoprecipitation and immunoblotting

Cells were lysed with lysis buffer consisting of 20 mM Tris-HCl (pH 7.4), 135 mM NaCl, 1% Triton X-100, 1% glycerol, and protease inhibitor cocktail tablets (Roche), incubated for 30 min at 4°C, and subjected to centrifugation at 14,000 g for 15 min at 4°C. The supernatants were boiled at 95°C for 5 min and then incubated with anti-FLAG, HA or Glu-Glu antibodies at 4°C for 90 min. After incubation with protein G-Sepharose 4B (GE Healthcare) at 4°C for 90 min, the beads were washed five times by lysis buffer and boiled at 95°C for 5 min. The proteins were resolved by SDS-PAGE (Novex gels; Invitrogen), transferred onto nitrocellulose membranes (iBlot; Life Technologies), blocked with PBS containing 0.05% Tween 20 and 5% skim milk, incubated with primary antibody at 4°C for 12 h, and then incubated with horseradish peroxidase (HRP)-conjugated secondary antibody at room temperature for 1 h. The immune complexes were visualized with Super Signal West Femto substrate (Pierce) and detected by an LAS-3000 image analyzer system (Fujifilm).

### Virus titration

Viral titers of HCV and JEV were determined by a focus forming assay as described previously (26). Viral titers of VSV and EMCV were quantified by a plaque forming assay using BHK-21 cells.

### Generation of USP15^-/-^ mice

USP15^-/-^ mice were generated as previously described using a C57BL/6N genetic background (41). The pX330 containing an sgRNA against the mouse *Usp15* gene was injected into mouse zygotes and transplanted into pseudopregnant female mice. The obtained mice (F1) were crossed with wild type mice and F2 mice and their DNA sequences were analyzed using the primers 5’-ATTTGGTACAGACCTGCCGG-3’ (forward) and 5’-TCGGAATAATGGGGAACTTGGG-3’ (reverse).

### Ethics Statement

All animal experiments were approved by the Institutional Committee of Laboratory Animal Experimentation (Research Institute for Microbial Diseases, Osaka University; project number: H27-06-0).

### VSV infection *in vivo*

USP15^+/+^, USP15^-/-^ and IFNα/ßR^-/-^ mice (16–17 weeks old) were intranasally infected with VSV (4 × 10^6^ pfu). Their survivals and body weights were monitored.

### Reporter assay

Huh7 cells, USP15KOHuh7 and USP15KOHuh7 cells expressing USP15 were transfected with pGL3-Basic SREBP-1c and pRL-TK and incubated for 2 days. Luciferase activity was determined by using a Dual-Luciferase Reporter Assay System (Promega).

### Treatment with palmitic acid and Oleic acid

Palmitic acid (PA) and oleic acid (OA) were dissolved in ethanol. Dissolved fatty acids were mixed with 10% fatty acid-free BSA (Sigma-Aldrich) to make a complex of fatty acid and BSA. Huh7 or USP15KOHuh7 cells were infected with HCV and treated with PA or OA for 4 days.

### Statistical analysis

All experiments were performed in triplicate with at least 3 independent experiments. All data represent the means ± SD of the independent experiments. The statistical analyses were performed using GraphPad Prism (GraphPad Prism Software). Significant differences were determined using Student’s *t*-test and are indicated with asterisks (*P<0.05) and double asterisks (**P<0.01) in each figure. Significant differences of *in vivo* survival data were determined using a log-rank test and are indicated with asterisks (*P<0.05) and double asterisks (**P<0.01) in each figure.

## Acknowledgements

The authors are grateful to M. Tomiyama for secretarial work, and M. Ishibashi for technical assistance. We also thank D.C.S. Huang, R. Bartenschlager, F. Chisari, and T. Wakita for providing experimental materials. The Core Instrumentation Facility of Osaka University conducted the FACS sorting.

This work was supported by the Program for Basic and Clinical Research on Hepatitis from the Japan Agency for Medical Research and Development (AMED) (JP18fk0210006h0003, JP17fk0210305h0003, JP18fk0210010h0003, JP18fk0210009h0503, JP17fk0210304h0003, JP18fk0210040h0001, and JP18fk0108036h0002), by the Ministry of Education, Culture, Sports, Science, and Technology (MEXT) of Japan (JP16H06432, JP16H06429, JP16K21723, JP15H04736, JP16K19139) and by the Research Program on Emerging and Re-emerging Infectious Diseases (17fk0108109h0001).

## Author contributions

T.O. and Y.M. designed the research; S.K., T.O. and Tatsuya Suzuki performed most of the experiments. Y.S., S.H., K.H., M.T., J.H., Z.H., D.V.C., H.I., P.D.N., Y.K., C.O. and T.F. assisted with the experiments; M.Y. provided the RNAi library; M.I. generated USP15^-/-^ mice; T.T., K.M., M.F., A.T. and Tetsuro Suzuki provided valuable materials; T. Satoh and S.A. provided IFNα/βR^-/-^ mice and assisted with the experiments; T.O. and Y.M. obtained research grants and wrote the manuscript.

